# Encoding of odor fear memories in the mouse olfactory cortex

**DOI:** 10.1101/297226

**Authors:** Claire Meissner-Bernard, Yulia Dembitskaya, Laurent Venance, Alexander Fleischmann

## Abstract

Odor memories are exceptionally robust and essential for animal survival. The olfactory (piriform) cortex has long been hypothesized to encode odor memories, yet the cellular substrates for olfactory learning and memory remain unknown. Here, using intersectional, *cFos*-based genetic manipulations (“Fos-tagging”), we show that olfactory fear conditioning activates sparse and distributed ensembles of neurons in mouse piriform cortex. We demonstrate that chemogenetic silencing of these Fos-tagged piriform ensembles selectively interferes with odor fear memory retrieval, but does not compromise basic odor detection and discrimination. Furthermore, chemogenetic reactivation of piriform neurons that were Fos-tagged during olfactory fear conditioning causes a decrease in exploratory behavior, mimicking odor-evoked fear memory recall. Together, our experiments identify odor-specific ensembles of piriform neurons as necessary and sufficient for odor fear memory recall.

## Introduction

Odor perception, and emotional and behavioral responses to odors strongly depend on experience, and learned odor-context associations often last for the lifetime of an animal (Mouly and Sullivan, 2010). The cellular and neural circuit mechanisms underlying olfactory learning and memory, however, remain poorly understood. Recent studies on episodic and contextual learning in hippocampal neural networks have suggested that memories are encoded in the activity of distributed ensembles of neurons, often referred to as a ‘memory trace’ (Mayford and Reijmers, 2015; Poo et al., 2016; Tonegawa et al., 2015). The neurons constituting such a memory trace are thought to encode information about the environmental context and associated emotions of past experiences, and their activity is necessary and sufficient for memory retrieval (Liu et al., 2012; Reijmers et al., 2007; Tanaka et al., 2014).

Here, we investigate the organization of odor memory traces in the olfactory (piriform) cortex of mice. The piriform cortex, a trilaminar paleocortical structure, is the largest cortical area receiving direct afferent inputs from the olfactory bulb, which, in turn, receives topographically organized inputs from olfactory sensory neurons in the nose. Individual piriform neurons respond to combinatorial inputs from multiple olfactory bulb projection neurons (Apicella et al., 2010; Davison and Ehlers, 2011), suggesting that odor objects are constructed in piriform cortex from the molecular features of odorants extracted in the periphery (Wilson and Sullivan, 2011). Recent electrophysiological and optical recordings have revealed that different odors activate distributed, overlapping ensembles of piriform neurons, which encode information about the identity and intensity of odors (Bolding and Franks, 2017; Miura et al., 2012; Roland et al., 2017; Stettler and Axel, 2009).

Piriform cortex has long been hypothesized to support auto-associative network functions that can retrieve previously learned information from partial or degraded sensory inputs (Haberly, 2001; Wilson and Sullivan, 2011). Piriform pyramidal cells form a large recurrent network, which is reciprocally connected with adjacent high-order associative areas including the prefrontal, entorhinal and perirhinal cortex and the amygdala (Johnson et al., 2000; Sadrian and Wilson, 2015). Storage of information is made possible by NMDA-dependent, associative plasticity of connections (Johenning et al., 2009; Kanter and Haberly, 1990; Quinlan et al., 2004). Furthermore, changes in piriform network activity (Chapuis and Wilson, 2011; Chen et al., 2011; Li et al., 2008; Sevelinges et al., 2004) and stabilization of piriform odor representations (Shakhawat et al., 2015) have been observed after associative olfactory learning. Finally, excitotoxic lesions of the posterior piriform cortex in rats perturb odor fear memories (Sacco and Sacchetti, 2010), and optogenetic stimulation of artificial piriform ensembles is sufficient to drive learned behaviors (Choi et al., 2011). Taken together, these studies have led to the hypothesis that piriform neural ensembles encode olfactory memory traces.

To test this hypothesis, we have developed an intersectional genetic strategy in mice to target piriform neurons that are activated by olfactory experience. We employed *cFos*-tTA transgenic mice (Garner et al., 2012), in which the tTA transcription factor is expressed under the promoter of the immediate early gene *cFos*, combined with virally targeted expression of tTA-responsive fluorescent reporters and regulators of neural activity (Alexander et al., 2009; Ferguson et al., 2011; Zhang et al., 2015). This approach allows us to selectively and persistently “Fos-tag” piriform neurons that were activated during olfactory fear conditioning. We then tested the behavioral consequences of manipulating such Fos-tagged piriform ensembles during memory recall. We find that chemogenetic silencing of piriform ensembles that were Fos-tagged during olfactory fear conditioning abolishes the expression of odor fear memories. In contrast, silencing piriform neurons responsive to a neutral odor only partially attenuates odor fear memory recall, suggesting that Fos-tagged piriform ensembles are largely odor-specific. Finally, artificial reactivation of Fos-tagged piriform neurons elicits behavioral changes consistent with memory recall. Together, our results suggest that piriform neural ensembles are an essential component of olfactory memory traces.

## Results

### Fos-tagging and functional manipulation of piriform neurons

To selectively label and manipulate piriform neurons that were activated during odor exposure, we used *cFos*-tTA transgenic mice (Reijmers et al., 2007), in which the activity-dependent *cFos* promoter drives expression of the tTA transcription factor, and we stereotaxically injected in their piriform cortex Adeno-Associated Viruses (AAVs) expressing Designer Receptors Exclusively Activated by Designer Drugs (DREADDs) under the control of the tTA-responsive promoter tetO (**Figure 1A**). DREADDs (Alexander et al., 2009; Ferguson et al., 2011) increase (hM3Dq:mCherry) or decrease (HA:hM4Di-IRES-mCitrine) the excitability of neurons upon activation by their ligand clozapine N-oxide (CNO). Temporal control of DREADD expression is provided by doxycycline, administered through the diet, which interferes with the binding of tTA to tetO and thus suppresses transgene expression (Gossen et al., 1995). Mice maintained in their home cage on a doxycycline-containing diet showed low basal DREADD expression, detected by anti-HA immunohistochemistry and native mCherry fluorescence (**Figure 1B and C**). To induce DREADD expression, mice were taken off doxycycline five days before odor presentation. Exposure to odor followed by foot shock (see **Experimental Procedures**) resulted in the Fos-tagging of sparse neural ensembles (median (interquartile range): 1.6 (0.3) % of piriform neurons, n=2 for hM4Di; 2.2 (0.5) % of piriform neurons, n=3 for hM3Dq), which were broadly dispersed throughout the anterior and posterior piriform cortex (**Figure 1B and C**), consistent with previous observations of endogenous cFos expression (Datiche et al., 2001; Illig and Haberly, 2003).

**Figure 1.**
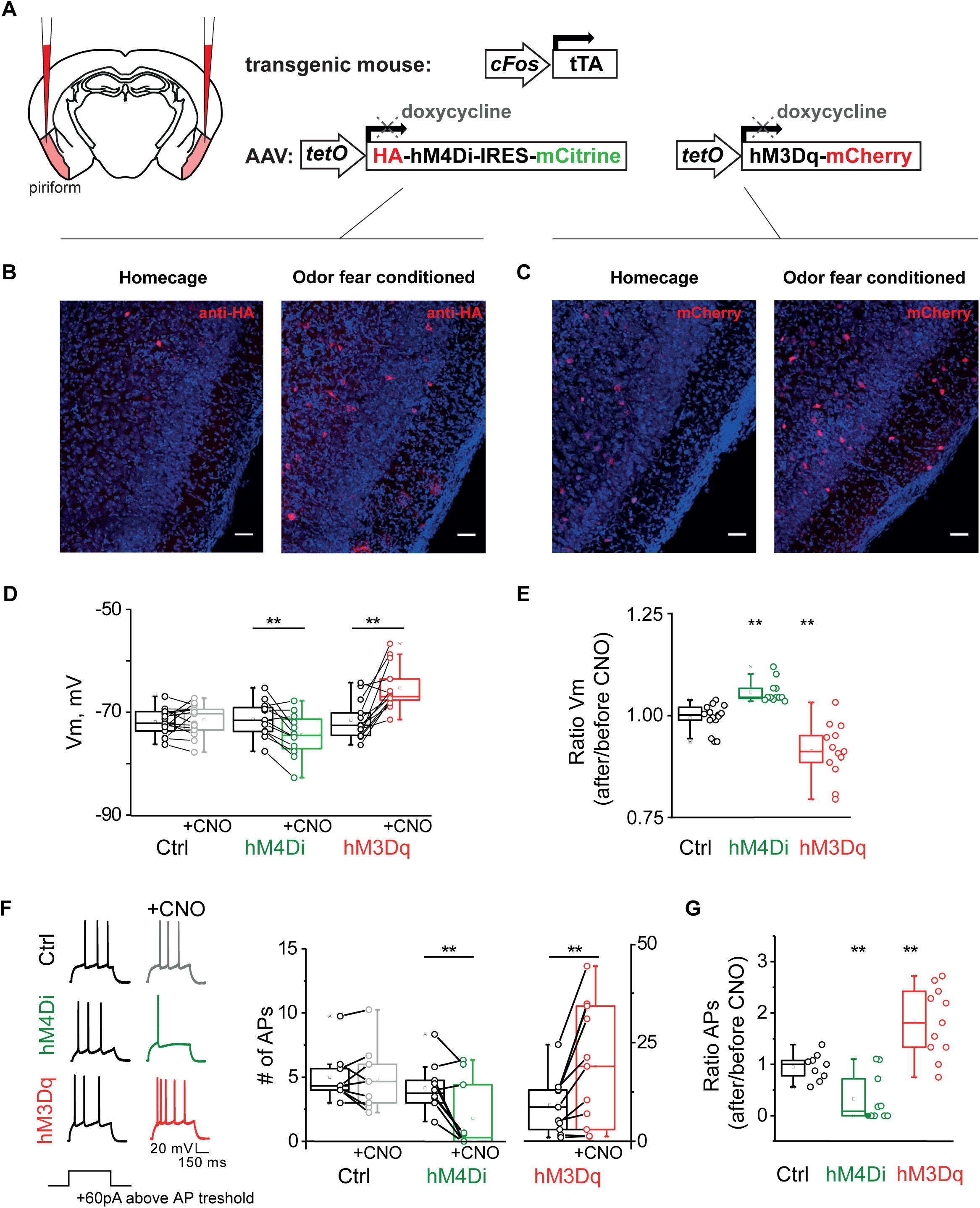
Fos-tagging and manipulation of piriform neurons. (A-C) Scheme of genetic strategy and histological characterization of Fos-tagged piriform ensembles. (A) The “silencing” and “activating” TetTag system. AAVs expressing DREADDs under the control of the tetO promoter (red) were injected in the piriform cortex of *cFos*-tTA transgenic mice. hM4Di or hM3Dq are expressed in active Fos-expressing piriform neurons upon doxycycline removal. (B-C) Mice remained in their homecage on doxycycline-containing diet (left), or were fear conditioned to ethyl-acetate five days after doxycycline removal (right). (B) Neurons expressing hM4Di were visualized using anti-HA immunohistochemistry. (C) Neurons expressing hM3Dq were visualized with mCherry fluorescence. Neurotrace counterstain in blue. Scale bars, 50 µm. (D-G) Electrophysiological characterization of Fos-tagged piriform neurons. (D) The resting membrane potential does not differ between DREADD-expressing and non-expressing cells. After CNO application, the resting membrane potential does not change in non-expressing cells (Ctrl), while hM4Di-expressing cells hyperpolarize and hM3Dq-expressing cells depolarize. (E) The relative change in the membrane potential after/before CNO does not change in non-expressing cells (Ctrl), while hM4Di-expressing cells hyperpolarize and hM3Dq-expressing cells depolarize. (F) The excitability of DREADD-expressing cells is measured by the number of action potentials (APs) resulting from the third depolarizing current step, counting from the first step that elicits spiking. Representative traces (left) and number of APs (right), before and after CNO application. After CNO application, the number of APs in DREADD-negative cells (Ctrl) does not change, significantly decreases in hM4Di-expressing cells and significantly increases in hM3Dq-expressing cells. (G) The relative change in the number of APs after/before CNO in DREADD-negative cells (Ctrl) does not change, significantly decreases in hM4Di-expressing cells and significantly increases in hM3Dq-expressing cells. Data are shown as individual data points, median and interquartile range, ** p<0.01.

To test whether CNO-mediated activation of DREADDs in Fos-tagged piriform neurons alters their excitability, we performed whole-cell recordings in acute brain slices containing piriform cortex. DREADD-expressing neurons were identified based on mCitrine (HA:hM4Di-IRES-mCitrine) or mCherry (hM3Dq:mCherry) fluorescence. Fos-tagged neurons displayed diverse morphologies and electrophysiological properties and included both excitatory and inhibitory neurons (**Supplementary Figure 1**). In the absence of the DREADD ligand CNO, the resting membrane potential of DREADD-expressing piriform neurons was indistinguishable from DREADD-negative cells (-71.70 (4.37) mV in DREADD-negative cells, n=24, -71.58 (4.32) mV in hM4Di-expressing cells, n=14, -72.56 (3.92) mV in hM3Dq-expressing cells, n=15, p=0.631, Kruskal-Wallis ANOVA, **Figure 1D**). After bath application of CNO (5 µM), the membrane properties selectively changed in DREADD-expressing cells: hM4Di-expressing cells hyperpolarized (-74.51 (5.17) mV, n=12, p=0.007, Wilcoxon Signed Ranks Test), and hM3Dq-expressing cells depolarized (-66.95 (4.12) mV, n=13, p=0.001, Wilcoxon Signed Ranks Test), whereas the resting membrane potential of DREADD-negative cells remained unchanged (-70.28 (3.81) mV, n=15, p=0.890, Wilcoxon Signed Ranks Test, **Figure 1D**). The relative changes were 1.00 (0.03) in DREADD-negative cells (n=15, p=0.890, One Sample Wilcoxon Signed Rank Test), 1.04 (0.03) in hM4Di-expressing cells, (n=12, p<0.001, One Sample Wilcoxon Signed Rank Test) and 0.91 (0.09) in hM3Dq-expressing cells, (n=13, p=0.001, One Sample Wilcoxon Signed Rank Test), **Figure 1E**). To determine the impact of CNO treatment on neuronal excitability, we delivered depolarizing current steps and compared the number of action potentials triggered by 60 pA step current above action potential threshold (see **Experimental Procedures**). We found that in hM4Di-expressing cells, the number of evoked action potentials significantly decreased after CNO application (3.75 (1.59) *vs*. 0.30 (3.48), n=10, p=0.010, Wilcoxon Signed Ranks Test), whereas in hM3Dq-expressing cells, the number of action potentials significantly increased after CNO application (8.67 (9.10) *vs*. 19.0 (27.6) mV, n=11, p=0.006, Wilcoxon Signed Ranks Test). In contrast, DREADD-negative cells did not exhibit a significant change in the number of evoked action potentials upon CNO application (4.33 (1.67) *vs*. 4.67 (3.00), n=9, p=0.672, Wilcoxon Signed Ranks Test, **Figure 1F**). The relative changes were 1.00 (0.41) in DREADD-negative cells, (n=9, p=0.641, One Sample Wilcoxon Signed Rank Tests), 0.09 (0.81) in hM4Di-expressing cells (n=10, p=0.010, One Sample Wilcoxon Signed Rank Tests) and 1.81±1.09 in hM3Dq-expressing cells (n=11, p=0.004, One Sample Wilcoxon Signed Rank Tests, **Figure 1G**). Taken together, these experiments show that Fos-tagging during olfactory fear conditioning marks a sparse and dispersed subpopulation of piriform neurons, and that CNO-mediated activation of hM4Di or hM3Dq expressed in Fos-tagged neurons selectively decreases or increases their excitability.

### Fos-tagged piriform ensembles are necessary for odor fear memory recall

If Fos-tagged piriform neurons that were activated during olfactory learning constitute an essential component of an olfactory memory trace, then perturbing the activity of these ensembles should interfere with memory recall. To test this prediction, we Fos-tagged piriform neurons during olfactory fear conditioning, and then chemogenetically silenced Fos-tagged ensembles during odor fear memory recall. *cFos*-tTA transgenic mice (“experimental group”, n=8) were bilaterally injected in piriform cortex with the AAV-tetO-hM4Di vector. Wild-type mice injected with the same virus served as a control (“control group”, n=8). During training, the conditioned odor (CS+) was presented at one of the ends of the conditioning box and paired with a foot shock, applied to the side of the box where the odor was delivered (see **Experimental Procedures** and **Figure 2A**). Mice thus learned to escape from the conditioned stimulus (CS+) by running towards the opposite side of the box. Two control odors, whose presentation was not paired with foot shock (CS-), were used to entrain odor-specific as opposed to generalized fear responses (Chen et al., 2011). Doxycycline was removed from the diet of mice prior to training, to permit the induction of hM4Di expression in active Fos-expressing piriform neurons of mice of the experimental group. Mice were returned to doxycycline-containing diet immediately after odor fear conditioning, and memory recall of the odor-foot shock association was tested three days later by presenting the odors alone in a new box (**Figure 2B and C**). All mice were intraperitoneally (i.p.) injected with CNO before testing, to exclude potential off-target effects by CNO (Gomez et al., 2017). For all mice, we verified viral gene expression by post hoc histological examination. Previous studies have suggested that the posterior piriform cortex is important for associative memory encoding (Sacco et al., 2010; Calu et al., 2008; Haberly 2001). We therefore excluded from our analysis mice in which parts of the posterior piriform cortex were spared from viral infection (**Supplementary Figure 2**). We combined two parameters to quantify the learned escape behavior: the position of the mouse in the box, and the increase in its maximum velocity after odor presentation (**Figure 2B**). These parameters were chosen because they described well the behavioral response of mice to the odor-foot shock pairing during training (**Supplementary Figure 3**).

**Figure 2.**
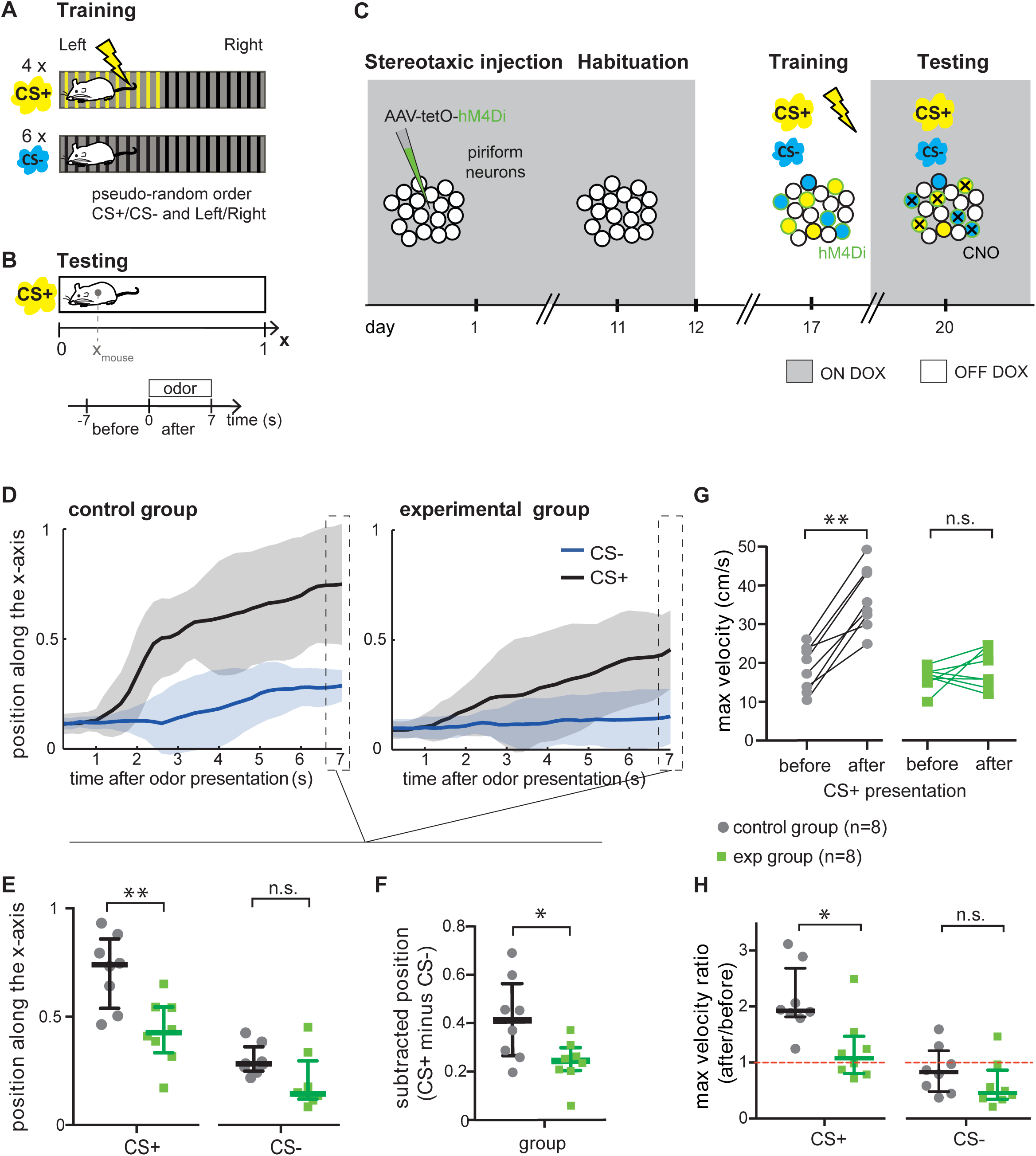
Fos-tagged piriform ensembles are necessary for odor fear memory retrieval. (A) During fear conditioning, mice are trained to associate ethyl-acetate (CS+) with foot shock. The CS+ is presented at one end of the training box, and the corresponding half-side of the box is electrified. Training includes the presentation of two other odors that are not paired with foot shock (CS-, eugenol and beta-citronellol). The CS+ is presented 4 times (2 times on each side) and each CS-three times. The order and side of odor presentations is pseudo-randomized. (B) During testing, the CS+ is presented 4 times (2 times on each side), and each CS-is presented once on each side, for 7 s each. The testing session is video-recorded and the position of the mouse’s centroid in the box along the x-axis is extracted (top). The length of the box is normalized to 1. Since the odor is presented randomly on the left or right side of the testing box, 0 is defined as the extremity where the odor is presented. For velocity measurements, a 7 s time window is defined both before and after stimulus onset (bottom). (C) Experimental design: *cFos*-tTA transgenic mice (experimental group, n=8) or wild-type mice (control group, n=8) are injected with AAV-tetO-hM4Di in both hemispheres of piriform cortex. Ten days later, mice are habituated to the training and testing boxes. Upon doxycycline (DOX) removal, mice are fear-conditioned, and hM4Di expression (green circles) is induced in Fos-tagged neurons (filled in blue and yellow). Mice are returned to DOX-containing diet to avoid further expression of hM4Di. Three days later, all mice are i.p. injected with CNO, which silences hM4Di-expressing neurons (black crosses), and memory retrieval is tested. (D to H) Olfactory memory retrieval after CNO injection, assessed by the position of mice in the testing box and their maximum velocity upon odor exposure. Memory retrieval was significantly impaired in *cFos*-tTA transgenic mice in which piriform neurons that were active during learning were chemogenetically silenced during memory recall (experimental group). (D) Position of mice in the testing box as a function of time after CS+ (black) and CS-(blue) presentation. (E) Position in the testing box 7 s after CS+ or CS-presentation. (F) Position 7 s after CS-presentation subtracted from the position 7 s after CS+ presentation. (G) Maximum velocity during a 7 s time window before and after CS+ presentation. (H) Maximum velocity ratio after/before CS+ (left) or CS-(right) presentation. Data are shown as median and interquartile range, * p<0.05; ** p<0.01; n.s. not significant.

As expected, mice in the control group exhibited robust escape behavior upon CS+ presentation. The median position of the mice in the box at the end of stimulus delivery was 0.74 (with 0 defined as the site of odor presentation and 1 as the opposite end of the box), and mice exhibited a 1.9-fold increase of their maximum velocity (**Figure 2E, G and H**). In contrast, escape behavior was significantly reduced in mice in which Fos-tagged neurons were silenced. Their median position in the box at the end of stimulus delivery, (0.42), and their 1.1-fold increase in maximum velocity were significantly lower than in the control group (Mann-Whitney test, U=7, p=0.007 for both position and velocity, **Figure 2E, G and H**). The difference in the position of mice after CS+ and CS-presentation was also significantly different between the control and experimental groups (Mann-Whitney test, U=13, p=0.0499, **Figure 2F**). Differences in behavioral responses of *cFos*-tTA transgenic mice were specifically due to hM4Di activation, as escape behavior after CNO injection was unaffected in *cFos*-tTA transgenic mice injected with AAVs expressing the calcium indicator GCaMP (n=9, **Supplementary Figure 4**).

All mice were similarly close to the odor port at the onset of CS+ odor presentation (Mann-Whitney test, U=28, p=0.7209, **Figure 2D**), and had similar maximum velocities during the 7 s time period that preceded odor presentation (Mann-Whitney test, U=24, p=0.4418, **Figure 2G**). These observations exclude any inherent differences between groups due to how the CS+ was presented. Furthermore, escape behavior in control mice was specific to the CS+, as none of the mice ran away from the CS-(**Figure 2D, E and H**). Finally, we did not observe any differences in the overall mobility of mice throughout the entire testing session. The median speed of mice of all groups (0.68 (0.28) cm/s and 0.46 (0.14) cm/s for the control and experimental groups respectively) was not significantly different (Mann-Whitney test, U=22, p=0.3282), nor did mice show a bias for the left or right side of the testing box (time spent on the left side divided by time spent on the right side: 1.36 (0.72) and 1.51 (0.52) for the control and experimental groups respectively, Mann-Whitney test, U=27, p=0.6454). Together, these data suggest that Fos-tagged piriform ensembles that were activated during olfactory fear conditioning are necessary for odor fear memory recall.

Even though *cFos*-tTA transgenic mice expressing hM4Di failed to robustly escape from the CS+, it should be noted that they behaved differently after CS+ and CS-presentation. They were further away from the odor port after CS+ presentation than after CS-presentation (Wilcoxon matched-pairs signed rank test, W=-36, p=0.0078, **Figure 2E**), and their maximum velocity ratio was significantly higher (Wilcoxon matched-pairs signed rank test, W=-36, p=0.0078, **Figure 2H**). This could indicate a partial memory of the learned association, possibly due to incomplete silencing of Fos-tagged neurons in piriform cortex, or the existence of parallel neural pathways that partially compensate for the loss of piriform functions. Finally, silencing of the piriform cortex in only one hemisphere did not abolish the learned escape behavior to the conditioned odor (**Supplementary Figure 5**).

### Silencing Fos-tagged piriform ensembles does not alter odor detection and discrimination

It is possible that the chemogenetic silencing of piriform ensembles that were Fos-tagged during olfactory fear conditioning disrupts odor detection and discrimination rather than selectively affecting odor fear memory recall. To test this possibility, we monitored sniffing behavior of mice during an olfactory habituation/dishabituation assay (**Figure 3A**), a well established test for odor detection and discrimination (Coronas-Samano et al., 2016; Verhagen et al., 2007; Wesson et al., 2008). Fos-tagging of piriform neurons during olfactory fear conditioning was performed as described above. Three days later, changes in sniffing behavior in response to odor exposure were tested in a plethysmograph, while hM4Di-expressing piriform neurons in *cFos*-tTA transgenic mice were silenced using i.p. injection of CNO (see **Experimental Procedures**). We observed that mice of both experimental (n=8) and control (n=6) groups increased their sniff frequency upon presentation of a novel, neutral odor (1.6 fold increase; Wilcoxon matched-pairs signed rank test, W=21, p=0.0312 and W=36, p=0.0078 for the control and experimental groups, respectively; **Figure 3B and C**). Repeated exposure to the same odor (neutral odor or CS-) resulted in a decrease in the sniff frequency, reflecting habituation after 4 consecutive exposures (**Figure 3C and D**). Finally, presentation of the CS+ increased sniff frequency (1.4 fold increase; Wilcoxon matched-pairs signed rank test, W=21, p=0.0312 and W=34, p=0.0156 for the control and experimental groups, respectively, **Figure 3D**). Such changes in odor sampling behavior have been shown to report the detection and discrimination of different odor stimuli (Coronas-Samano et al., 2016; Verhagen et al., 2007; Wesson et al., 2008). Therefore, these data suggest that basic behaviors characteristic of odor sampling, detection and discrimination were unaffected by the silencing of piriform ensembles that were Fos-tagged during olfactory fear conditioning.

**Figure 3.**
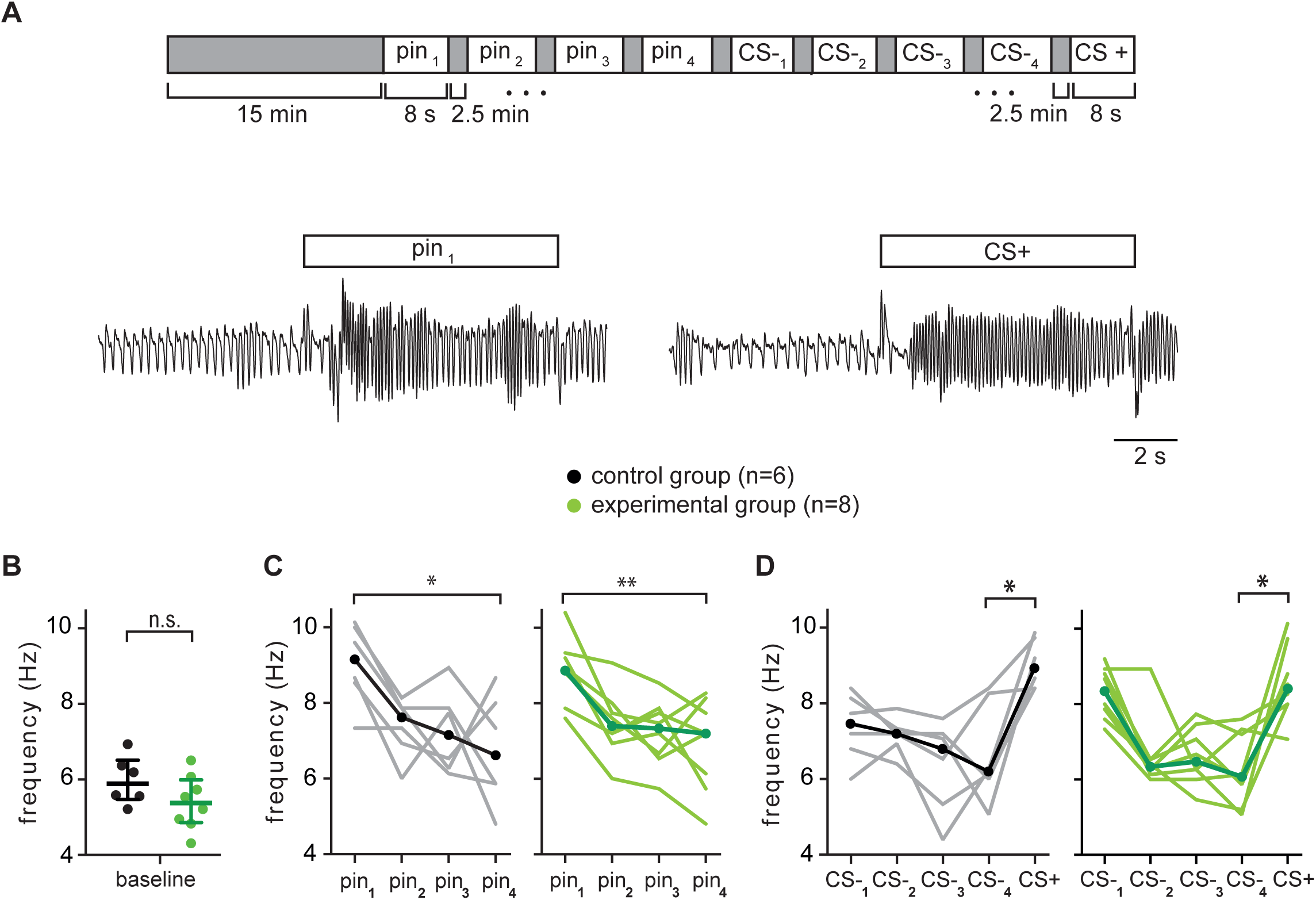
Silencing Fos-tagged piriform ensembles does not interfere with odor detection and discrimination. (A) (Top) Experimental design: Fos-tagging during olfactory fear conditioning is performed as described in Figure 2. Three days after fear conditioning, sniffing behavior is quantified in a plethysmograph. Mice are i.p. injected with CNO, habituated to the plethysmograph for 15 min, and each odor is then presented for 8 s with a 2.5 min inter-trial interval. pin: pinene, CS-: beta-citronellol, CS+: ethyl-acetate. (Bottom) Representative sniff recordings from one mouse when exposed to pinene or the CS+ (ethyl-acetate). (B) Baseline sniff frequency, corresponding to the averaged sniff frequency 8 s before odor onset. (C-D) For both the control (n=6) and experimental group (n=8), repeated exposure to the same odor resulted in a decrease in the sniff frequency (habituation), while presentation of a different odor resulted in an increase in the sniff frequency (dishabituation). (C) Sniff frequency for the first to fourth presentation of pinene: control (left) and experimental group (right). (D) Sniff frequency for the first to fourth presentation of the CS-, and subsequent presentation of the CS+: control (left) and experimental group (right). Each thin line represents data of individual mice, the circles and thick lines represent averaged data. Data are shown as median and interquartile range, * p<0.05; ** p<0.01; n.s.: not significant

### Fos-tagged piriform ensembles are odor-specific

Odors activate unique but overlapping ensembles of piriform neurons (Bolding and Franks, 2017; Poo and Isaacson, 2009; Roland et al., 2017; Stettler and Axel, 2009). Therefore, silencing neurons that respond to one odor could partially interfere with the retrieval of information associated with other odors. To test this prediction we modified the behavioral protocol to separate Fos-tagging from olfactory fear conditioning. *cFos*-tTA transgenic and wild-type control mice were injected with AAV-tetO-hM4Di, and expression of hM4Di in piriform neurons was induced while mice were exposed to odor (eugenol or beta-citronellol) in a neutral environment, without subsequent foot shock. Fos-tagging resulted in the labeling of sparse neural ensembles, similar to those labeled during odor-foot shock association (1.1 (0.3) % of piriform neurons, n=3). Mice were returned to doxycycline-containing diet and trained two days later to associate a different odor (ethyl-acetate, CS+) with foot shock. Behavioral testing of the learned escape behavior in response to the CS+ was performed one day later, after i.p. injection of CNO (see **Experimental Procedures** and **Figure 4A**). We found that learned escape behavior of *cFos*-tTA transgenic mice expressing hM4Di (experimental group, n=12) was similar but somewhat attenuated compared to wild-type controls (n=10). Both groups exhibited an escape behavior after CS+ presentation, indicated by the significant increase in the maximum velocity (Wilcoxon matched-pairs signed rank test, W=55, p=0.0020 and W=56, p=0.0269 for the control and experimental groups respectively; no difference in the maximum velocity ratio after/before between groups, Mann-Whitney test, U=35, p=0.1071, **Figure 4E and F**). The difference in the position of mice after CS+ and CS-presentation was also similar between the control and experimental groups (Mann-Whitney test, U=37, p=0.1402, **Figure 4D**). However, we observed that silencing neutral odor representations somewhat dampened the behavioral response to the conditioned stimulus, as behavioral responses were less robust compared to controls when quantifying the position of mice in the box after CS+ presentation (Mann-Whitney test, U=22, p=0.0112, **Figure 4B and C**). The attenuated behavioral responses we observe likely reflect the partial degradation of odor information as a result of the silencing of piriform neurons responsive to both the CS+ and the neutral odor.

**Figure 4.**
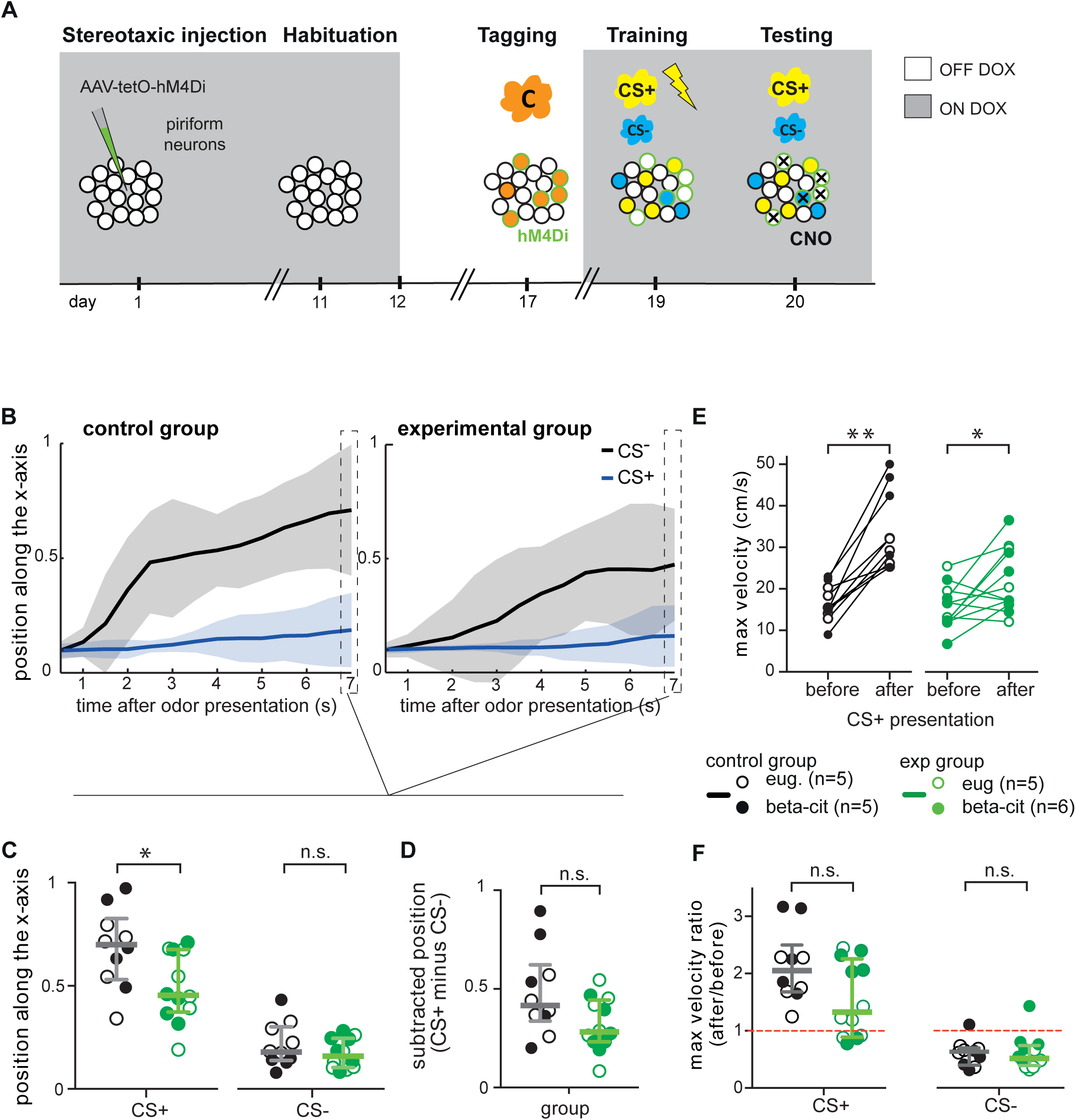
Fos-tagged piriform ensembles are odor-specific. (A) Experimental design: dissociating Fos-tagging and fear conditioning. Upon doxycycline (DOX) removal, mice are presented with an odor in a neutral environment (eugenol, eug. or beta-citronellol, beta-cit.). hM4Di expression (green circles) is induced in Fos-tagged neurons (filled in orange). Two days later, mice are fear conditioned to ethyl-acetate (CS-: limonene and beta-cit. when Fos-tagged with eug., limonene and eug. when Fos-tagged with beta-cit.). One day later, memory retrieval is tested in the presence of CNO, which silences hM4Di-expressing neurons (black crosses). (B-F) Olfactory memory retrieval after CNO injection, assessed by the position of mice in the testing box and their maximum velocity. Memory retrieval was moderately attenuated in *cFos*-tTA mice expressing hM4Di in neurons that were Fos-tagged during presentation of a neutral odor (experimental group, n=12), compared to WT mice (control group, n=10). (B) Position of mice as a function of time after CS+ and CS-presentation. (C) Position of mice in the testing box 7 s after CS+ or CS-presentation. (D) Position 7 s after CS-presentation subtracted to the position 7 s after CS+ presentation. (E) Maximum velocity during the 7 s before and after CS+ presentation. (F) Maximum velocity ratio after/before CS+ (left) or CS-(right) presentation. Data are shown as median and interquartile range, * p<0.05; ** p<0.01; n.s. not significant.

### Reactivation of Fos-tagged piriform ensembles is sufficient to retrieve an odor fear memory

If piriform neural ensembles that were activated during olfactory fear conditioning encode an odor fear memory trace, then reactivation of these neurons may be sufficient to trigger memory recall. To test this prediction, we devised a modified behavioral task to assess memory recall. We monitored exploratory behavior in an open field during ambient odor exposure or artificial reactivation of Fos-tagged neurons, a test that does not rely on an escape behavior from an odor originating from a spatially defined odor source (**Figure 5A**). We first trained mice to associate ethyl-acetate, the CS+, with foot shock. Upon ambient exposure to the CS+, diffused above the open field, exploratory behavior of mice was significantly attenuated compared to baseline exploratory behavior measured one day earlier (Wilcoxon matched-pairs signed rank test, n=6, W=-21, p=0.0312). Ethyl-acetate exposure did not affect exploratory behavior of mice that had previously been exposed to ethyl-acetate without foot shock (Wilcoxon matched-pairs signed rank test, n=5, W=1, p>0.9999). These data suggest that re-exposure of mice to the CS+ causes an increase in anxiety-like behavior, expressed as a decrease in exploratory behavior in the open field.

**Figure 5.**
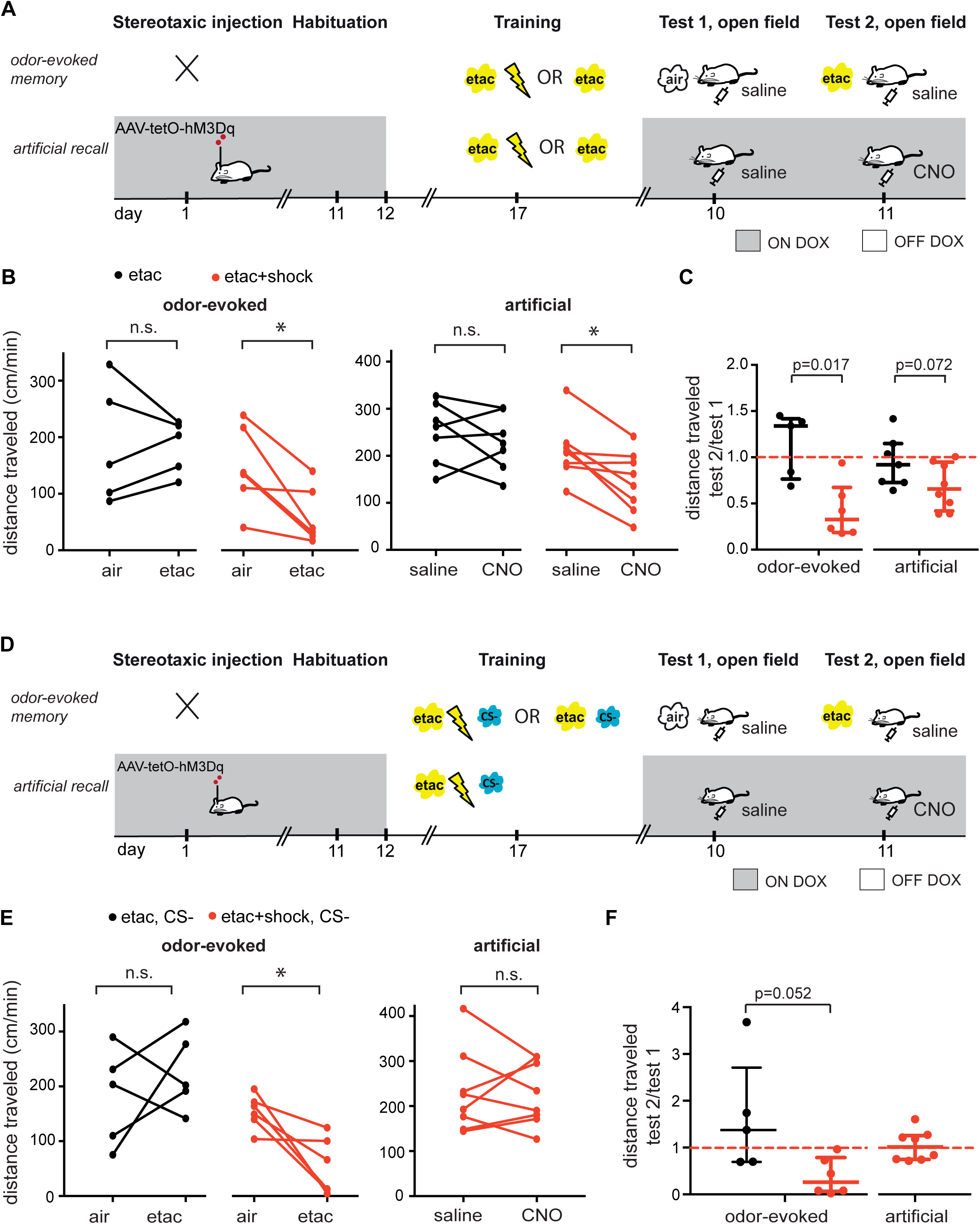
Reactivation of Fos-tagged piriform ensembles is sufficient for odor fear memory recall. (A) Experimental design: odor-evoked and artificial odor fear memory recall tested in an open field assay. Mice are trained to associate ethyl-acetate with foot shock (CS+), and mice exposed to ethyl-acetate without foot shock serve as controls. Wild-type mice are used for testing odor-evoked fear memory recall. For artificial memory recall, *cFos*-tTA transgenic mice are injected with AAV-tetO-hM3Dq in piriform cortex to induce hM3Dq expression in Fos-expressing neurons during training. Two days later, baseline exploratory activity in an open field is measured during 22 min, 5 min after i.p. injection of saline (test 1). The next day (test 2), activity of mice in the open field is measured again. For odor-evoked memory recall, ethyl-acetate is presented above the open field from minutes 15 to 18 of the test 2 session. For artificial memory recall, CNO is i.p injected to reactivate Fos-tagged, hM3Dq-expressing piriform neurons. (B) As a measure for exploratory activity, the distance travelled is computed from minute 15 to 18 (corresponding to odor exposure during test 2). For artificial memory recall, the distance travelled is computed over the entire testing session. Ethyl-acetate exposure and CNO injection lead to a decrease in exploratory behavior in mice previously fear conditioned to ethyl-acetate (n=6 and n=8 respectively). Previous exposure to ethyl-acetate without foot shock does not alter exploratory behavior upon re-exposure to ethyl-acetate or re-activation of Fos-tagged piriform ensembles (n=5 and n=7 respectively). (C) Distance traveled ratio test 2/test 1. (D) Experimental design: same as in (A), except that training includes the presentation of two other odors that are not paired with foot shock (CS-, eugenol and beta-citronellol). (E) Ethyl-acetate exposure leads to a decrease in exploratory behavior in mice previously fear-conditioned to ethyl-acetate (n=6). However, reactivation of piriform ensembles Fos-tagged during olfactory fear conditioning including reinforced and non-reinforced odor stimuli does not affect exploratory behavior (n=8). (F) Distance traveled ratio test 2/test 1. Data are presented as median and interquartile range. * p<0.05; n.s. not significant.

We next repeated the experiment in *cFos*-tTA transgenic mice, in which the piriform cortex had been infected with the activating DREADD AAV-tetO-hM3Dq (**Supplementary Figure 6**). Instead of exposure to the CS+, we i.p. injected the DREADD ligand CNO to reactivate hM3Dq-expressing piriform neurons that were Fos-tagged during odor-foot shock association (see **Experimental Procedures** and **Figure 5A**). We found that CNO-mediated reactivation, similar to CS+ exposure, caused a significant decrease in exploratory behavior of mice that had undergone olfactory fear conditioning (Wilcoxon matched-pairs signed rank test, n=8, W=-34, p=0.0156, **Figure 5B and C**). CNO-mediated reactivation of piriform neurons that were Fos-tagged during ethyl-acetate exposure without foot shock did not affect exploratory behavior of mice (Wilcoxon matched-pairs signed rank test, n=7, W=-10, p=0.4688, **Figure 5B and C**). Taken together, these data suggest that chemogenetic reactivation of piriform neurons that were active during odor-foot shock exposure is sufficient to trigger fear memory recall.

We next asked whether artificial memory recall depends on the specificity of the Fos-tagged neural ensemble: is piriform reactivation sufficient to trigger fear memory recall, as long as it includes the CS+-tagged ensemble? For this purpose, we generated a synthetic Fos-tagged ensemble, by sequentially exposing mice to ethyl-acetate paired with foot-shock (CS+) and neutral odors (CS-, eugenol and beta-citronellol) (**Figure 5D**). While exposure to the CS+ resulted in anxiety-like behavior similar to what was observed in the previous experiment (Wilcoxon matched-pairs signed rank test, n=6, W=-21, p=0.0312, **Figure 5E and F**), artificial reactivation of synthetic Fos-tagged ensemble did not result in changes in exploratory behavior that would indicate memory recall (Wilcoxon matched-pairs signed rank test, n=8, W=-4, p=0.8438, **Figure 5E and F**). These data suggest that during artificial reactivation, piriform cortex cannot extract meaningful information from the synthetic representation generated by the sequential presentation of CS+ and CS-odors.

## Discussion

The piriform cortex has long been thought to provide the substrate for storing associative olfactory memories. Several studies have shown that piriform cortex cells can encode information that carries behavioral significance (Calu et al., 2008, Mandairon et al., 2014, Sacco et al., 2010), yet their functional relevance has never been assessed by direct experimental manipulation. Using a *cFos*-dependent, intersectional genetic approach to visualize and manipulate piriform neurons activated during olfactory fear conditioning (**Figure 1**), we found that chemogenetic silencing of Fos-tagged piriform ensembles abolished a learned odor escape behavior (**Figure 2**). Silencing these Fos-tagged ensembles did not alter basic odor detection and discrimination (**Figure 3**). Furthermore, chemogenetic silencing of neurons responsive to a neutral odor only moderately attenuated odor fear memory recall, suggesting that Fos-tagged piriform ensembles are largely odor-specific (**Figure 4**). Finally, specific chemogenetic reactivation of piriform ensembles that were active during learning resulted in reduced exploratory behavior in an open field assay, an effect that was similarly observed when exposing mice to the conditioned odor stimulus (**Figure 5**). Together, our experiments identify piriform neurons that were active during learning as necessary and sufficient to trigger odor fear memory recall. This indicates the formation of functional connections between Fos-tagged piriform cells and other associative areas and firmly establish piriform ensembles as an essential component of odor fear memory traces.

### Artificial reactivation of an olfactory memory trace

Reactivation of neurons by CNO-mediated activation of DREADD receptors does not recapitulate the temporal characteristics of piriform odor responses and their modulation by active sampling (Bolding and Franks, 2017; Miura et al., 2012). Despite this, reactivation of Fos-tagged piriform ensembles activated during odor-foot shock exposure was sufficient to elicit a behavioral response. How piriform neural circuits and downstream target regions process such artificial activity patterns remains to be determined. One possibility is that despite the temporal limitations of memory trace reactivation, piriform network mechanisms can retrieve the perception of the reinforced conditioned stimulus. Consistent with this model, recent studies have shown that spatial patterns of odor-evoked activity, in the absence of precise temporal information, are sufficient to decode odor identity in piriform cortex (Bolding and Franks, 2017; Miura et al., 2012; Roland et al., 2017). Alternatively, it is possible that reactivation generates a state of stress and anxiety, but without evoking the perception of the odor. Candidate target areas for the processing of odor-fear associations include the basolateral amygdala and the medial prefrontal cortex (Dejean et al., 2016; Gore et al., 2015), however, the relevant neural circuit components remain to be identified. Finally, reactivating a synthetic ensemble of neurons activated during exposure to the reinforced stimulus (CS+) and the non-reinforced stimuli (CS-) did not elicit a measurable behavioral response. This observation suggests that behaviorally relevant information cannot be extracted from a neural ensemble representing conflicting (aversive versus neutral) information. This result is consistent with the finding that the reactivation of an artificial contextual memory in the hippocampus competes with the retrieval of a learned context-shock association (Ramirez et al., 2013).

### Limitations of Fos-tagging for the study of olfactory memory traces

A general limitation of the c*Fos*-tTA system is that the tagging of active neurons is transient. The duration of the expression of tTA-dependent proteins is determined by the time course of induction and the stability of *cFos*-tTA-dependent transcripts and proteins in tagged neurons. Previous studies have shown that TetTag-dependent expression of regulators of neural activity lasts for at least five days but becomes undetectable by thirty days (Cowansage et al., 2014; Liu et al., 2012; Zhang et al., 2015). We therefore restricted our manipulations of neural activity and behavioral analyses to a three-day time window after Fos-tagging of neurons during olfactory conditioning. Another constraint of the Fos-tagging system is its slow temporal dynamics. As a consequence, neurons responding to the reinforced conditioned stimulus (CS+), the odorant paired with foot shock, and neurons responding to the non-reinforced conditioned stimuli (CS-), the odorants not associated with foot shock, were tagged in our silencing experiments. The use of a CS-compromises the specificity of the Fos-tagged neural ensembles. Nevertheless, the CS-provides an important experimental readout for the odor selectivity of the behavioral response.

Interestingly, the number of Fos-tagged piriform neurons we observe is significantly lower than the number of odor-responsive neurons detected in electrophysiological recordings in awake mice (Bolding and Franks, 2017; Miura et al., 2012; Zhan and Luo, 2010). However, in these experiments, a large fraction of cells exhibited low firing rates. Therefore, the Fos-tagged population could represent a subpopulation of neurons that is strongly activated by odor (Schoenenberger et al., 2009). Fos-tagging could also mark plastic changes supporting memory formation (Cole et al., 1989; Minatohara et al., 2015). Indeed, *cFos* mRNA levels decrease significantly after injection of an antagonist of the NMDA receptor, a key player in the induction of synaptic plasticity (Tayler et al., 2011), and long-term memory and synaptic plasticity are impaired when cFos production is perturbed in the central nervous system (Fleischmann et al., 2003; Seoane et al., 2012). Future studies will allow to directly test these hypotheses.

### Memory traces in hippocampus and piriform cortex

The hippocampus has been studied extensively for its role in spatial and contextual memory (Basu and Siegelbaum, 2015). Recently, Fos-tagging of hippocampal neurons during contextual fear conditioning has provided important insight into the cellular and neural circuit mechanisms of learning and memory (Cai et al., 2016; Garner et al., 2012; Liu et al., 2012; Ramirez et al., 2013; Roy et al., 2017; Ryan et al., 2015). However, if principles of memory formation and storage in hippocampus-related neural networks apply to other cortical structures remains largely unknown. Interestingly, pirform cortex and hippocampus share a similar circuit organization and both have been modeled as auto-associative networks (Haberly, 2001). Consistent with this model, our findings reveal striking similarities between olfactory memory traces in the piriform cortex and memory traces of contextual fear in the hippocampus. In both piriform cortex and hippocampus, learning activates sparse and distributed ensembles of neurons that appear to lack topographic organization. Furthermore, hippocampal neurons tagged during contextual fear conditioning, and piriform neurons tagged during olfactory fear conditioning were both necessary and sufficient for memory retrieval (Liu et al., 2012; Tanaka et al., 2014), The olfactory system is particularly well-suited for studying learning and memory. Indeed, odor memories and olfactory-driven behaviors are robust in animals, including in mice, providing highly reliable and quantitative behavioral readouts. Furthermore, the olfactory cortex is only two synapses away from the peripheral olfactory sensory neurons in the olfactory epithelium, and the neural inputs that drive piriform cortex activity have been well characterized (Banerjee et al., 2015; Dhawale et al., 2010; Economo et al., 2016; Fukunaga et al., 2012; Kato et al., 2012; Roland et al., 2016; Yamada et al., 2017). Knowing the properties of neural input patterns will provide the basis for understanding neural circuit input-output computations and their modification by learning and experience.

## Acknowledgements

We thank Mark Mayford for sharing the *cFos*-tTA transgenic mouse line and Brian Roth and Susumu Tonegawa for sharing plasmids. We thank Yves Dupraz and Gérard Parésys for their work on the behavioral apparatuses, Agatha Anet for help with behavioral experiments, Tristan Piolot for help with image acquisition, Philippe Mailly for help with cell quantification and Marion Ruinart de Brimont for help with cloning. We thank Karim Benchenane and Sophie Bagur for sharing their plethysmograph. We thank Anne Didier, Kevin Franks, Boris Gourévitch, Andreas Schaefer, and Claire Zhang for careful reading and critical comments of the manuscript. This work was supported by the “Amorçage de jeunes équipes” program (AJE201106) of the Fondation pour la Recherche Médicale (to A.F.), and a postdoctoral fellowship by the LabEx “MemoLife” (to C.M.B.).

## Author contributions

Conceptualization, CMB and A.F.; Methodology, investigation and analysis, C.M.B, Y.D. (electrophysiology); Writing of original draft, C.M.B and A.F., with contributions from Y.D. and L.V.; Supervision and funding acquisition, C.M.B., L.V. and A.F.

## Material and Methods

### Mice

Mice were housed at 24°C with a 12-hour light/12-hour dark cycle with standard food and water provided ad libitum. Mice were group-housed with littermates until the beginning of surgery and then single-housed in ventilated cages throughout the duration of the experiment. *cFos*-tTA mice (Reijmers et al., 2007) were fed a diet containing 1g/kg doxycycline for a minimum of 4 days before surgery. Wild-type control animals were siblings that did not carry the *cFos*-tTA transgene. The age of mice at the time of behavioral testing ranged from 10 to 15 weeks. Male mice were used for the hM4Di-mediated neural silencing, and females for the hM3Dq-mediated neural activation experiments. Experiments were carried out according to European and French national institutional animal care guidelines (protocol APAFIS#2016012909576100).

### Constructs and viruses

The hSynapsin promoter of pAAV-hSyn-HA-hM4Di-IRES-mCitrine (kindly provided by Dr. B. Roth, University of North Carolina at Chapel Hill) was excised using the restriction enzyme XbaI (New England Biolabs) and replaced by the Tetracycline Response Element of a pAAV-TRE-EYFP (excised with Xba1 et Nhe1, plasmid kindly provided by Dr. S. Tonegawa, MIT, Cambridge). The pAAV-pTRE-tight-hM3Dq-mCherry (Zhang et al., 2015) was purchased from Addgene (**#**66795). Adeno-associated viruses (AAVs) were generated at Penn Vector Core, University of Pennsylvania (serotype 8, 10^13^ genome copies/mL, 1:2 dilution with sterile PBS on the day of injection).

### Stereotaxic injection

Mice were anaesthetized with ketamine/xylazine (100 mg/kg/10 mg/kg, Sigma-Aldrich) and AAV vectors were injected stereotaxically into the piriform cortex (coordinates relative to bregma: anterior-posterior, -0.6 mm; dorsal-ventral, -4.05 mm; lateral, 4.05 mm and -4.05 mm). Using a micromanipulator and injection assembly kit (Narishige; WPI), a pulled glass micropipette (Dutscher, 075054) was slowly lowered into the brain and left for 30 seconds in place before infusion of the virus at an injection rate of 0.2-0.3 μL per min. 0.8-1 μL of virus was sufficient to infect a large area of the piriform cortex. The micropipette was left in place for an additional 3-4 min and then slowly withdrawn to minimize diffusion along the injector tract.

### Immunohistochemistry

Mice were deeply anesthetized with pentobarbital and perfused transcardially with phosphate-buffered saline (PBS) followed by 4% paraformaldehyde. Brains were post-fixed 4h in 4% paraformaldehyde, and 100-200 µm coronal sections were cut with a vibrating blade microtome (Microm Microtech). Sections were permeabilized in 0.1% Triton-X100/PBS (PBST) for 1 h, then blocked in 2% heat-inactivated horse serum (HIHS)-PBST for 1 h. Sections were incubated in 2% HIHS-PBST containing polyclonal primary antibodies (rabbit anti-HA, 1:200, Cell Signaling; rabbit anti-GFP, 1:1000, Invitrogen) with gentle agitation at 4°C overnight. Next, sections were rinsed 3 times in PBST for 20 min and incubated in NeuroTrace 640/660 (N21483, Invitrogen) and species-appropriate Alexa Fluor 488 (green) and Alexa Fluor 568 (red)-conjugated secondary antibodies (1:1000, Invitrogen) at 4°C overnight. Sections were washed 2 times in PBST and 1 time in PBS, mounted on slides and coverslipped with Vectashield mounting medium (Vectorlabs). Images were acquired as Z-stacks (70 to 140 μm in total thickness, step size 7 μm) with a Leica SP5 confocal microscope or as single plane sections with a Zeiss Axio Zoom microscope and processed in Fiji.

### Counting

Z-stacks (7 µm step) were acquired with a Leica SP5 confocal microscope with a 20X objective and a resolution of 512×512 or 1024×1024 pixels (1 pixel: 0:72µm). First, the piriform cortex was delineated on a maximum projection of each stack using a custom-written ImageJ macro. For each mouse, 2 to 4 sections (median volume 9.3e-2 (9.4e-3) mm^3^) were analyzed, from one or both hemispheres, at Y=-0.8 and Y=-1.6mm relative to bregma.

HA-stained neurons (hM4Di) were counted by hand. hM3Dq-mCherry positive neurons were counted using a custom-written ImageJ plugin. After pre-processing the image (Subtract Background, Remove Outliers and Median Filter), the z-stack is thresholded using the RenyiEntropy algorithm. On each slice of the stack, objects with an area smaller than 40µm^2^ are removed using the Analyze Particle command in ImageJ. A mask of each slice from the stack is created containing the filled outline of the measured particles. The number of cells is the number of 3D objects detected in the stack. Automated cell detection can be prone to error, thus, the detection efficiency was checked on a few pictures by two people blind to the experimental conditions.

The numbers of cells were calculated as numbers of cells per cubic millimeter. An estimate of the total number of neurons (counterstained with NeuroTrace) per cubic millimeter (7.1e4 mm^−3^) was obtained by counting neurons in representative volumes containing the three layers of piriform cortex. Numbers are consistent with those obtained by (Srinivasan and Stevens, 2017).

### Electrophysiology

Parasagittal or coronal slices (300 µm thick) of piriform cortex were prepared from 6-8 week-old *cFos*-tTA mice injected with AAV-tetO-hM4Di-IRES-mCitrine or AAV-tetO-hM3Dq-mCherry. Animals were anaesthetized with ketamine and xylazine (100 mg/kg/10 mg/kg, Sigma-Aldrich), perfused with ice-cold ACSF (125 mM NaCl, 2.5 mM KCl, 25 mM glucose 25mM NaHCO_3_, 1.25 mM NaH_2_PO_4_, 2 mM CaCl_2_, 1 mM MgCl_2_, 1 mM pyruvic acid, bubbled with 95% O_2_ and 5% CO_2_ and adjusted to 295±5 mOsm osmolarity), and decapitated. The brain was cooled with ice-cold ACSF solution and then sliced using a 7000SM2 vibrating microtome (Campden Instruments, UK). Slices were incubated in the same solution at 34°C for one hour and then continuously perfused with ACSF solution (2 mL/min) at 34°C in the recording chamber. DREADD expression was detected with two-photon excitation (830nm, ChameleonMRU-X1, Coherent, UK) under a Scientifica TriM Scope II microscope (LaVision, Germany), with a 60x water-immersion objective. Whole-cell recording pipettes with 5-7 MΩ resistance were filled with the following solution (in mM): 122 Kgluconate, 13 KCl, 10 HEPES, 10 phosphocreatine, 4 Mg-ATP, 0.3 Na-GTP, 0.3 EGTA (adjusted to pH 7.35 with KOH). The recording solution also contained the morphological tracer Alexa Fluor 594 (5 µM, red channel) for non-expressing and hM4Di-positive cells or the Ca^**2**+^-sensitive dye Fura-2 (300 µM, replacing EGTA in the recording solution, green channel) for hM3Dq-positive cells, to enable identification of the patched cell and to visualize its morphology. The excitability of cells was measured in current-clamp mode by 500 ms steps of current injections from -300 to +500 pA with steps of 20 pA. We compared the number of action potentials triggered by equivalent depolarization at each step 60 pA above the action potential threshold before and after CNO (5 µM) application. The series resistance was usually <20 MΩ, and data were discarded if it changed by more than 20% during the recording. Signals were amplified using EPC10-2 amplifiers (HEKA Elektronik, Lambrecht, Germany). Voltage-clamp recordings were filtered at 5 kHz and sampled at 10 kHz, and current-clamp recordings were filtered at 10 kHz and sampled at 20 kHz, with the Patchmaster v2×32 program (HEKA Elektronik).

### Behavioral apparatus

A training box was used to train mice to escape from an odor. The box was rectangular (L 57 cm, W 17 cm, H 64 cm), with a grid floor made of 72 stainless-steel rods (diameter=6 mm, space between rods=2 mm). Current was delivered by an aversive stimulator (MedAssociates, 115V, 60 Hz). A custom-made switcher allowed an electric foot shock (0.6 mA, 0.6 ms) to be applied independently to either half of the box. A testing box was used to test memory retrieval. Its dimensions were similar to the training box. Materials for the walls and floor were different from the training box to create a different context. The open field box consisted of a white Plexiglas (6 mm thickness) container (50 cm x 50 cm x 38 height).

Sniffing behavior was monitored in freely moving mice using a plethysmograph (Emka Technologies). Electrical signals were amplified using RHD 2132 amplifier boards (Intan Technologies) and band-pass filtered (1-30 Hz).

Odors were delivered using an 8 channels olfactometer (Automate Scientific). Continuous air flowed through a bottle of mineral oil (Sigma-Aldrich) to avoid pressure changes when presenting the odors. The following odors (Sigma-Aldrich), at a concentration of 1% (vol/vol) in mineral oil were used: ethyl-acetate, beta-citronellol, eugenol, limonene, and pinene.

### Behavioral procedures

AAVs were stereotaxically injected in both hemispheres of piriform cortex of *cFos*-tTA transgenic mice and littermate wild-type controls. Mice were given 10 days to recover, before being habituated on day 11 to the training and testing/open field/plethysmograph boxes (40 and 10 min each). 24 hours after habituation, doxycycline was removed and replaced by a normal diet. Five days later, mice were fear conditioned to ethyl-acetate or exposed to an odor (odor “C”) in a neutral environment. Immediately after fear conditioning or odor presentation, mice were put back on a diet containing doxycycline. During fear conditioning, ethyl-acetate (the CS+) was presented at one extremity of the training box, and the corresponding half-side of the box’ floor was electrified for 0.6 s (0.6 mA). The foot shock was delivered 4 seconds after CS+ presentation, and the CS+ lasted for an additional 2 seconds. Mice learned to escape to the opposite side when the CS+ was presented. The CS+ was presented 4 times, 2 times on each side of the box. When fear conditioning included presentation of two non-reinforced conditioned stimuli (CS-1 and CS-2), the choice between CS+ and CS-as well as the side of presentation was pseudo-randomized. Training always started with one presentation of each CS-followed by the CS+. The two CS-odors were presented during 7 seconds, 3 times each, with a total presentation of 3 on the right side and 3 on the left side. During testing, the CS+ was presented 4 times (2 times on each side), and each CS- was presented once on each side, for 7 s each. As for the training session, the order and side of odor presentations was pseudo-randomized: testing always started with one presentation of each CS-, followed by one CS+ presentation.

### Drugs

Clozapine-N-oxide (C0832, Sigma-Aldrich) was first dissolved in dimethylsulfoxide (DMSO, D2438, Sigma, final concentration of 1% vol/vol), then further diluted in 0.9**%** sterile saline solution to a final concentration of 0.2 mg/mL. Aliquots were stored for up to two months at -20°C and equilibrated to room temperature before injection. The solution was administered intraperitoneally (3 mg/kg). After injection, mice were left undisturbed for 25 minutes (Figure 2 and 4) or 5 minutes (Figure 3 and 5) in their home cage before the start of the experiment.

### Data analysis

Electrophysiological data were analyzed with custom-made software in Python 3.0 and averaged in MS Excel (Microsoft, USA). Behavioral sessions were video recorded and analyzed using custom-written MatLab programs (available upon request). Parameters extracted from multiple CS+ (CS-) presentation were averaged for each mouse.

### Statistics

Statistical analysis was performed with Prism (GraphPad). Non-parametric tests were used: Mann-Whitney test for between group comparison, and Wilcoxon matched-pairs signed rank test for within group comparison. Values are represented as median (interquartile range) unless otherwise stated. The n represents number of mice, or number of recorded neurons for the electrophysiology experiment.

